# Selection-, age-, and exercise-dependence of skeletal muscle gene expression patterns in a rat model of metabolic fitness

**DOI:** 10.1101/013706

**Authors:** Yu-yu Ren, Lauren G. Koch, Steven L. Britton, Nathan R. Qi, Mary K. Treutelaar, Charles F. Burant, Jun Z. Li

**Author notes:** Corresponding Author: Jun Z. Li, Ph.D., Department of Human Genetics, 5940A Buhl, Box 5618, University of Michigan, Ann Arbor, MI 48109.

## Abstract

Aerobic exercise capacity can influence many complex traits including obesity and type 2 diabetes. We established two rat lines by divergent selection of intrinsic aerobic capacity. The high capacity runners (HCR) and low capacity runners (LCR) differed by ~9-fold in aerobic capacity after 32 generations, and diverged in body fat, blood glucose, and other health indicators. To study the interplay among genetic differentiation, age, and strenuous exercise, we performed microarray-based gene expression analyses in skeletal muscle with a 2×2×2 design to compare HCR and LCR, old and young animals, and between rest and exhaustion, for a total of eight groups (n=6 each). Transcripts for mitochondrial function are expressed higher in HCR than LCR at both rest and exhaustion, for both age groups. Extracellular matrix components decrease with age in both lines and both rest and exhaustion. Interestingly, age-effects in many pathways are more pronounced in LCR, suggesting that HCR’s higher innate aerobic capacity underlies both increased lifespan and heathspan.

## Introduction

Aerobic capacity in humans is strongly associated with longevity and a reduction in a variety of traits associated with poor metabolic health. To study the interplay between intrinsic aerobic capacity and metabolic fitness, we selectively bred rats for maximal running capacity to generate two strains which also resulted in divergence for a variety of metabolic trait [1]. The two lines, termed high capacity runners (HCR) and low capacity runners (LCR), originated from a founder population of genetically heterogeneous rats originally derived from outcrossing 8 inbred strains (N:NIH stock) [2]. The animals were not trained, so the differences in fitness were due to physiological differences acquired through the selective breeding. Through this selective breeding, there is a divergence for a number of phenotypic traits with the LCR demonstrating higher levels of triglyceride, free fatty acids, glucose and insulin levels and a higher body weight and adiposity [3, 4]. HCR rats show higher levels of maximal oxygen consumption, skeletal muscle oxidative enzyme levels and mitochondrial content [5-7]. We have recently shown that an increased capacity for skeletal muscle fatty acid and branched chain amino acid oxidation underlies the higher oxidative capacity in these animals (Overmyer et al., in revision). As in humans, the enhanced oxidative capacity of the HCR compared to the LCR is paralleled by increases in lifespan, with the median age of death increasing from 23.5 months for LCR to 30.1 months for HCR rats, representing a 28% difference in life expectancy, with no significant difference for maximal lifespan between females and males within lines.

In as early as Generation-7 of selection, HCR displayed significantly greater O_2_ utilization in the skeletal muscles [5]. Continued selective breeding up to generation 15 resulted in further divergence in O_2_ utilization as well as O_2_ delivery in the skeletal muscle [8]. An initial gene expression analysis of the skeletal muscle from HCR and LCR at generation 18 revealed significant differences for genes in the pathways of oxidative energy metabolism, including fat metabolism, branched-chain amino acid metabolism, Krebs cycle, and oxidative phosphorylation [9]. A subsequent study that looked at gene expression in skeletal muscle of HCR and LCR at generation 16 found that HCR upregulated genes involved in lipid metabolism and fatty acid elongation compared to LCR in exercise-trained rats, while the sedentary rats only showed minor differences in gene expression between the two lines [7]. The differences in gene expression were found to be consistent with results from proteomic analysis of skeletal muscle mitochondria (Overmyer et al., In Revision) which showed similar pathways enriched in HCR vs. HCR.

It has long been appreciated that biological regulation, in this case transcript levels, are affected by inherited genetic variation, naturally occurring aging process, as well as responses to immediate physiological stressors. These factors often act jointly but have not been analyzed simultaneously in a single study. Here we analyzed the transcriptomic profiles of both young and aged female rats from generations 29 and 32, and under both resting and exercise conditions with the goal of identifying pathways that could explain the divergence in aerobic capacity, longevity, and adaptation to exercise.

## Results

Our study adopted a 2×2×2 full factorial design to simultaneously examine the effects of three factors: HCR-LCR, aging (Old-Young), exercise (Rest-Exhaustion), and their interactions. Here “Young” animals are of age 31.1 ± 2.6 weeks and “Old” are 99.4 ± 2.9 weeks. Among the young animals, HCRs and LCRs show 8.5-fold difference in their maximal running capacity. Compared to the young, running performance in old animals declined by ~50% in both lines, and the HCR-LCR difference remained at 8.9-fold (**Supplementary Figure 1**). Within each age group in either line there was not a significant correlation between age and maximal running distance (**Supplementary Figure 1**). For three factors and two levels each (see **Methods**), there are eight experimental combinations. In each combination we measured the skeletal muscle (extensor digitorum longus, EDL) samples of six animals as biological replicates. Tissues from “Exhaust” animals were obtained immediately (<10 mins) after the run-to-exhaustion test. In all, we measured 19,607 transcripts in 48 samples in a single batch of microarray experiments.

### Global patterns

A principal component analysis (PCA) of the 19,607 measured transcripts separates the 48 samples into eight clusters in the PC1-PC2 space; and they correspond to the eight known groups, as marked by the colored ellipsoids (**Figure 1**). The eight clusters occupy mostly non-overlapping areas in the PC1-PC2 space. While a few of the clusters are close to each other, most are “coherent” and have gaps of varying sizes to the nearest cluster. Thus, at the global level there are observable transcriptional effects for all three factors. PC1 is mainly driven by the Old-Young differences, while both PC2 and PC3 are driven by HCR-LCR and exercise effects (**Figure 1**). PC3 is driven by the difference between the HCR-Rest animals (for both Old and Young) and the other six groups (**Supplementary Figure 2**).

**Figure 1.**
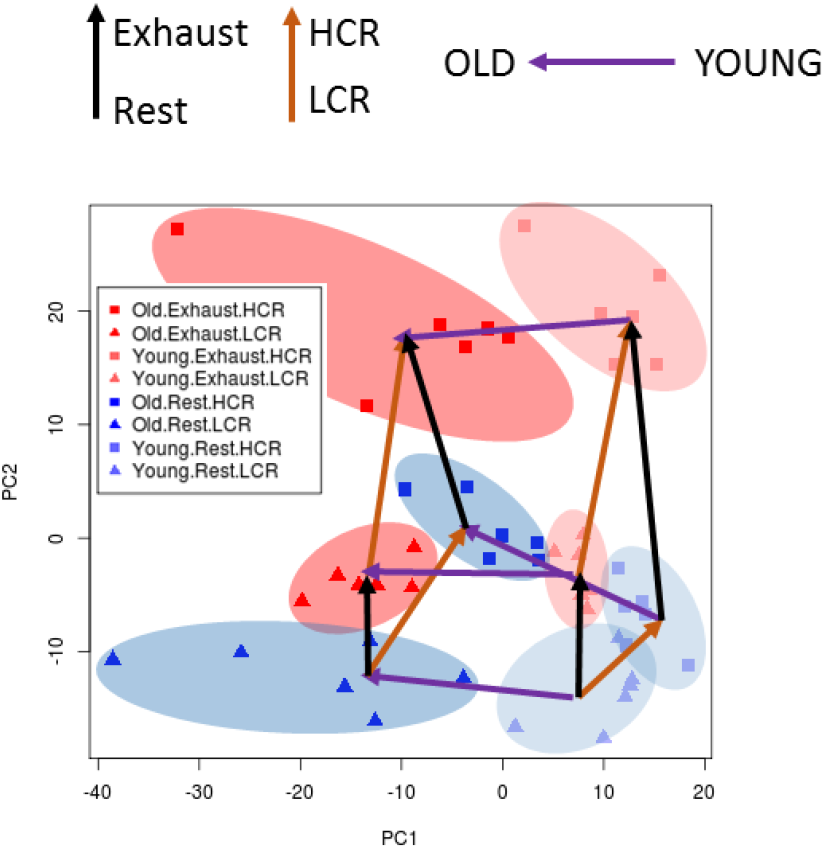
shown is a principal component analysis (PCA) plot (PC1 vs PC2) for 48 rats across the expression of ~20K transcripts. The samples are divided across 8 different groups in a 2×2×2 design to compare HCR and LCR, aged and young, and between rest and exhaustion (n=6 each). Groups are highlighted using different colored ovals to show clustering, and HCR-LCR are shown as squares and triangles, respectively. All of the exhausted animals are red, and animals at rest are blue. Old animals are shown as dark red/blue, and young animals are shown as light red/blue. Arrows are drawn from each cluster to show the direction of each group variable; purple arrows show direction of old-young animals; orange arrows show direction of HCR-LCR; black arrows show direction of exhaust-rest.

As a 2×2×2 design can be naturally displayed as a cube, we overlaid a hexahedron, i.e., an irregular, “stretched” cube, in **Figure 1** to connect the median expression patterns of the eight groups. Note that this hexahedron is not a quantitative representation of the high-dimension gene expression profiles, nor the between-cluster distances in their reduced two-dimensional view, but only a geographic illustration of the relationships among the experiment groups. This representation has six quadrilateral faces (or planes) forming three opposing pairs, each representing the two levels of a given factor. For example, the left and right faces represent the Old and Young animals, respectively, whereas the top and bottom faces represent Exhaust and Rest groups, respectively. The stretched cube has eight vertices, representing the centroid of the eight experimental groups; and its twelve edges represent the twelve two-way contrasts, each representing the main effect of a given factor in one of the four strata formed by the other two factors. In **Figure 2**, we used a three-letter shorthand to indicate the eight vertices, where “H” and “L” denote “HCR” and “LCR”, respectively, “O” and “Y” denote “Old” and “Young”, respectively, and “R” and “E” denote “Rest” and “Exhaust”, respectively. For example, the upper left vertex of the front face, “L-O-E”, is for the Old, LCR animals measured at Exhaustion. The set of four quasiparallel edges connecting a pair of opposing faces are shown as arrows of the same color: red for HCR-LCR, purple for Old-Young, and black for Rest-Exhaust. The twelve edges were also numbered 1-12 for ease of description. For example, Edge-2, from L-O-E to H-O-E, is the HCR-to-LCR difference for Old and Exhausted animals. This contrast is also written as (HCR-LCR|Old, Exhausted) in **Figure 3**, where “|” is the mathematical notation of “conditional on”, indicating the specific strata in which the contrast in defined.

**Figure 2.**
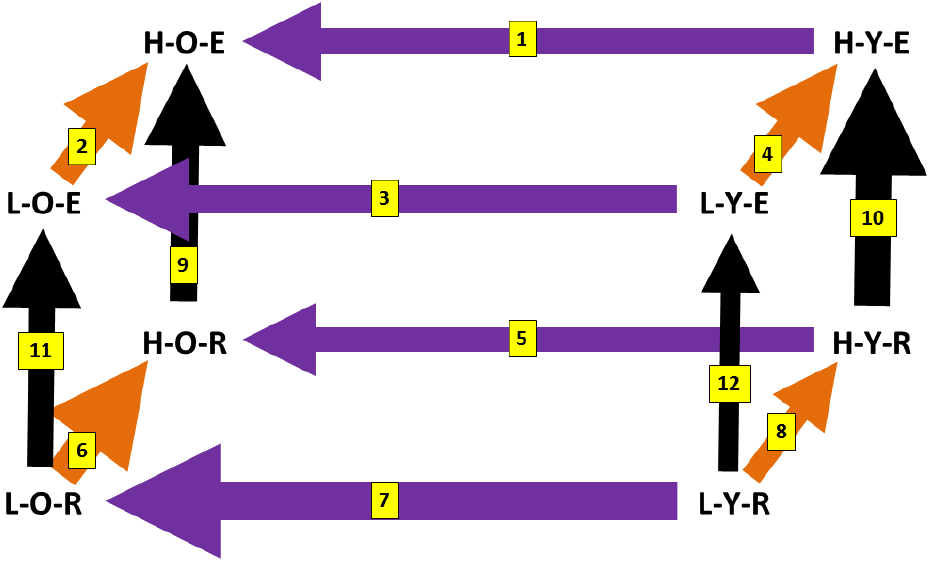
shown is a cube depiction of the PCA plot in Figure 1. Each group is represented by their abbreviated names (HCR/LCR = H/L, old/young = O/Y, exhaust/rest = E/R). The arrow color scheme is the same as Figure 1. (A) The numbers beside each arrow represents the Euclidean distance calculated using all ~20K transcripts, with larger numbers representing more distant groups in terms of skeletal muscle transcriptome. (B) The numbers on each arrow represent the denotation of the analysis (between 1-12) that will be referred to in the manuscript, and the thickness of each arrow represents the Euclidean distance for each side.

**Figure 3.**
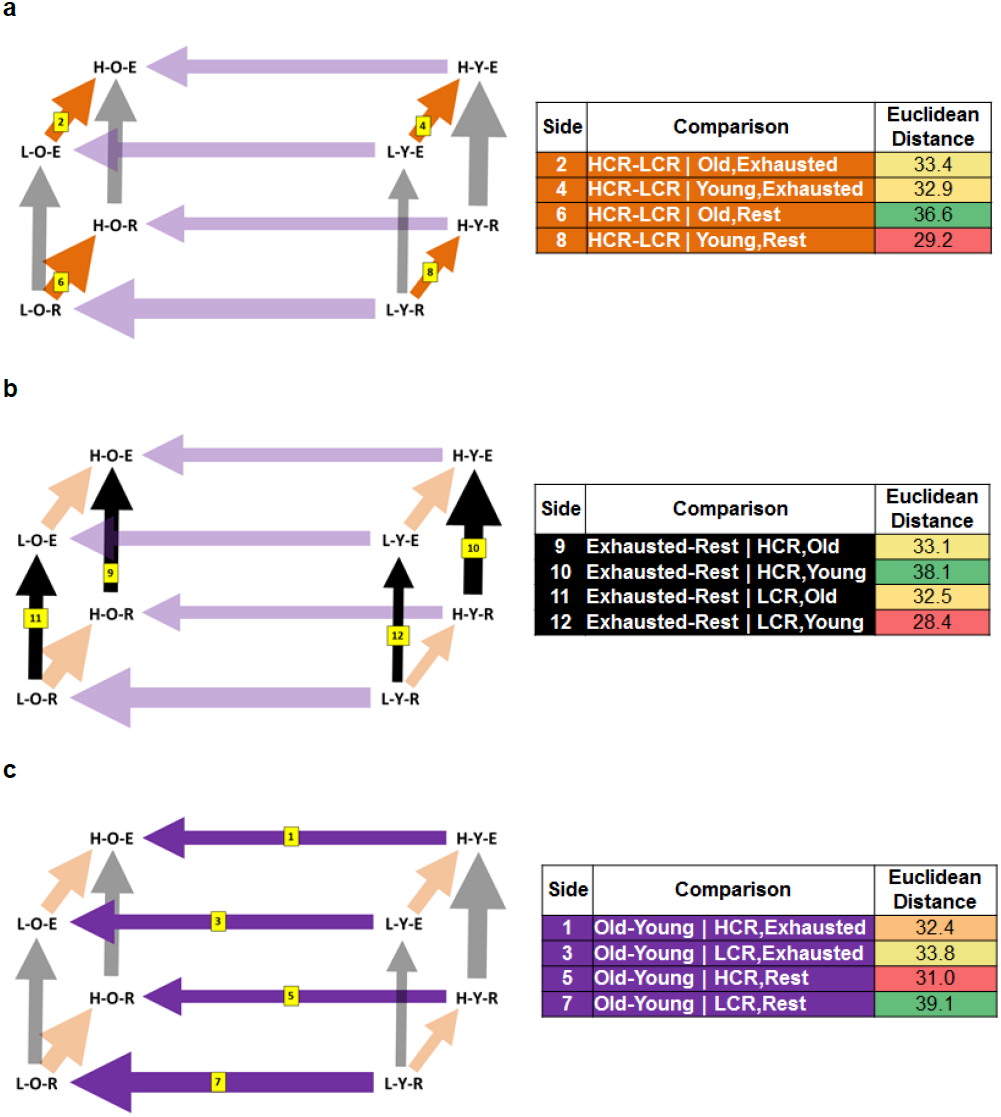
(A) shown are the cube depiction and Euclidean distances focused on the effects of genetic background: differences between HCR and LCR (sides 2, 4, 6, and 8); (B) shown are the cube depiction and Euclidean distances focused on the effects of exercise: differences between exhaustion and rest (sides 9, 10, 11, and 12); (C) shown are the cube depiction and Euclidean distances focused on the effects of aging: old vs young (sides 1, 3, 5, and 7).

Globally, the eight clusters form a well-proportioned convex cube, indicating that each of the three factors has a main effect, that the effects are comparable among the line, age, and exercise factors, and that they jointly determine the observed gene expression pattern. Further, the effects are not strictly additive (or independent). If they were, i.e., if there was no interaction among the factors, all six faces would be parallelograms, i.e., formed by parallel edges of equal lengths, and the opposing faces would form parallel pairs of planes. This this were the case, with suitable rotation of the PC axes the six faces could all be transformed to rectangles. However, the observed hexahedron is not a cuboid: it contains unparalleled faces and unparalleled edges; and in most faces, the opposing edges are of unequal length, indicating that the effect of any one factor depends on the specific combination of the levels of the other factors: the classic definition of statistical interaction. Our simultaneous analyses of the three factors thus revealed both main effects and their interactions, as examined in more details below.

We calculated the genomic distance along the twelve edges using all 19,607 transcripts, and in the cube display, varied the line widths to be proportional to the genomic distance (**Figure 2**). This way, a thicker line indicates a larger contrast (or distance), thus providing a different, but analogous, visual representation as the stretched cube shown in **Figure 1**, where it was the line lengths that represent the effect size along individual edges. **Figure 2** used the Euclidean distance as line width; the numeric values were shown in **Figures 3**. Alternative distance measures, such as the median absolute difference (MAD) between pairs of group centroids, yield similar results: the Pearson’s correlation coefficient (r) between MAD and Euclidean values is 0.91 across the 12 conditions. In the following we will describe the analysis of the three factors, one at a time, before describing the analysis of two-factor interactions.

### Between-line differences (HCR vs. LCR)

For each gene we assessed the HCR-LCR main effect overall, corresponding to the transcriptomic differences between the two genetic lines, averaged over Old-Young and Rest-Exhaust conditions. In the geographic representation this corresponds to the distance between the center of the HCR face (the back plane of the cube) to the center of the LCR face (front plane). The effect size reflects the transcriptomic consequence after divergent selection for aerobic running capacity. In all, 2,838 transcripts are significantly difference at Benjamini-Hochberg False Discovery Rate (BH-FDR) < 0.05 (**Supplementary Table 1**). The pathway analysis of these genes will be described in a later section.

We next analyzed the HCR-LCR difference separately for each of the four age-exercise combinations. Across the four strata, Old-Rest has the largest line contrasts, while Young-Rest has the smallest. This can be interpreted from two perspectives. First, *the HCR-LCR effect at Rest is age-dependent*: the lengths of Edge-6 and Edge-8, defined as Euclidean distance over all measured genes, are 36.6 and 29.2, respectively, indicating a stronger between-line difference in the Old animals (**Figure 3a**). In contrast to Rest, this age dependence of line effect is much reduced at Exhaustion: the lengths of Edge-4 and Edge-2 are 32.9 and 33.4, respectively, nearly the same between the Old and Young animals. From the second but equivalent perspective, *the HCR-LCR difference for Young animals depends on the exercise state*: it is greater at Exhaustion (Edge-4 vs. Edge-8; 32.9 vs. 29.2), but conversely, for Old animals the HCR-LCR difference is greater at Rest (Edge-2 vs. Edge-6; 33.4 vs. 36.6).

### Exercise effects (Exhaustion vs. Rest)

Next, we assessed the Exhausted-Rest main effect averaged over the HCR-LCR and Old-Young conditions. This corresponds to the distance between the center of the Exhausted face (top of the cube) to the center of the Rest face (bottom of the cube). The effect size reflects the transcriptomic adaptation after an endurance run. In all, 1,715 transcripts are significantly difference at BH-FDR < 0.05 (**Supplementary Table 1**). The pathway analysis of these genes will be described below.

We then analyzed the Exhausted-Rest difference separately for each of the four line-age combinations. Across the four strata, HCR-Young has the largest line contrasts whereas LCR-Young has the smallest. At Young age, the Exhausted-Rest effect is line-dependent: the lengths of Edge-10 and Edge-12 are 38.0 and 28.4, respectively, indicating a stronger exercise difference in the HCR animals (**Figure 3b**), which may reflect the longer exercise-related stimulus in the HCR due to the enhanced running capacity in the HCRs compared to LCRs. *This line-dependence of exercise effect is much reduced when measured in Old animals:* the lengths of Edge-9 and Edge-11 are 33.1 and 32.5, respectively. In an alternative view, *the age-dependence of exercise effect varies by line*: in HCR it is greater for Young animals (Edge-10 vs. Edge-9; 38.0 vs. 33.1); but conversely, in LCR the exercise difference is greater for Old animals (Edge-11 vs. Edge-12; 32.5 vs. 28.4).

### Aging effects (Old vs. Young)

The Old-Young main effect, averaged over HCR-LCR and Exhausted-Rest conditions, corresponds to the distance between the center of the Old face (left plane of the cube) to the center of the Young face (right plane). The effect size reflects the transcriptomic changes during the aging process. In all, 2,561 genes are significantly difference at BH-FDR < 0.05 (**Supplementary Table 1**). The pathway analysis of these genes will be described below.

Again we analyzed the Old-Young difference separately for each of the four line-exercise combinations. Across the four strata, LCR-Rest has the largest line contrasts whereas HCR-Rest has the smallest. At Rest, *the Old-Young effect is line-dependent*: the lengths of Edge-5 and Edge-7 are 31.0 and 39.1, respectively, indicating a stronger age difference in the LCR animals (**Figure 3c**). This line dependence of age effect is much reduced when measured at Exhaustion: the lengths of Edge-1 and Edge-3 are 32.4 and 33.8, respectively. In an alternative view, the Old-Young difference for HCR is slightly greater for Exhausted animals (Edge-1 vs. Edge-5; 32.4 vs. 31.0), but conversely, for LCR the age difference is greater for Rest animals (Edge-3 vs. Edge-7; 33.8 vs. 39.1).

### Pathway analyses of the three factors

The three overall comparisons, for the main effects of line, age, and exercise, respectively, implicated many biological pathways (**Supplementary Table 2**), of which we focus on five most strongly affected. These non-overlapping Gene Ontology pathways are: *Mitochondria Part, Extracellular Matrix, Collagen Fibril Organization, Focal Adhesion,* and *Sequence-Specific DNA Binding Transcription Factor Activity* (**Table 1**). Stratified analysis of each of the three effects in the four combinations of the other two factors, as shown by the twelve edges in Figure 1, showed largely consistent patterns as the overall effects (**Table 2** **and** **Figure 4**).

**Table 1.** shown are the -log(p-values) for the major pathway groups in the overall main effects of the three factors. A heatmap is used to show the difference in significance levels; with green denoting greater significance and red denoting less significance.

| Analysis | Comparison | Mitochondrial Part | Extracellular Matrix | Collagen Fibril Organization | Focal Adhesion | Sequence-Specific DNA Binding Transcription Factor Activity |
| --- | --- | --- | --- | --- | --- | --- |
| Line | HCR-LCR | 65.9 | 0.90 | 1.6 | 0.18 | 0.76 |
| Age | Old-Young | 0.07 | 55.1 | 25.0 | 16.9 | 6.9 |
| Exercise | Exhaust-Rest | 6.5 | 3.2 | 0.10 | 1.0 | 27.6 |

**Table 2.**
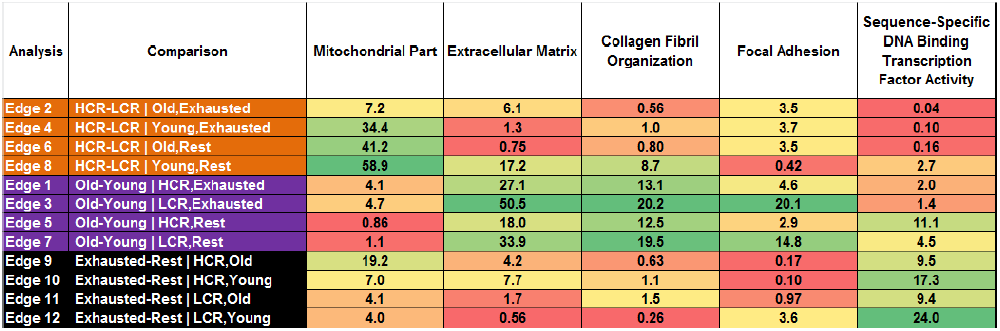
shown are the -log(p-values) for the major pathway groups in each main effect analysis. A heatmap is used to show the difference in significance levels; with green denoting greater significance and red denoting less significance.

**Table 3.**
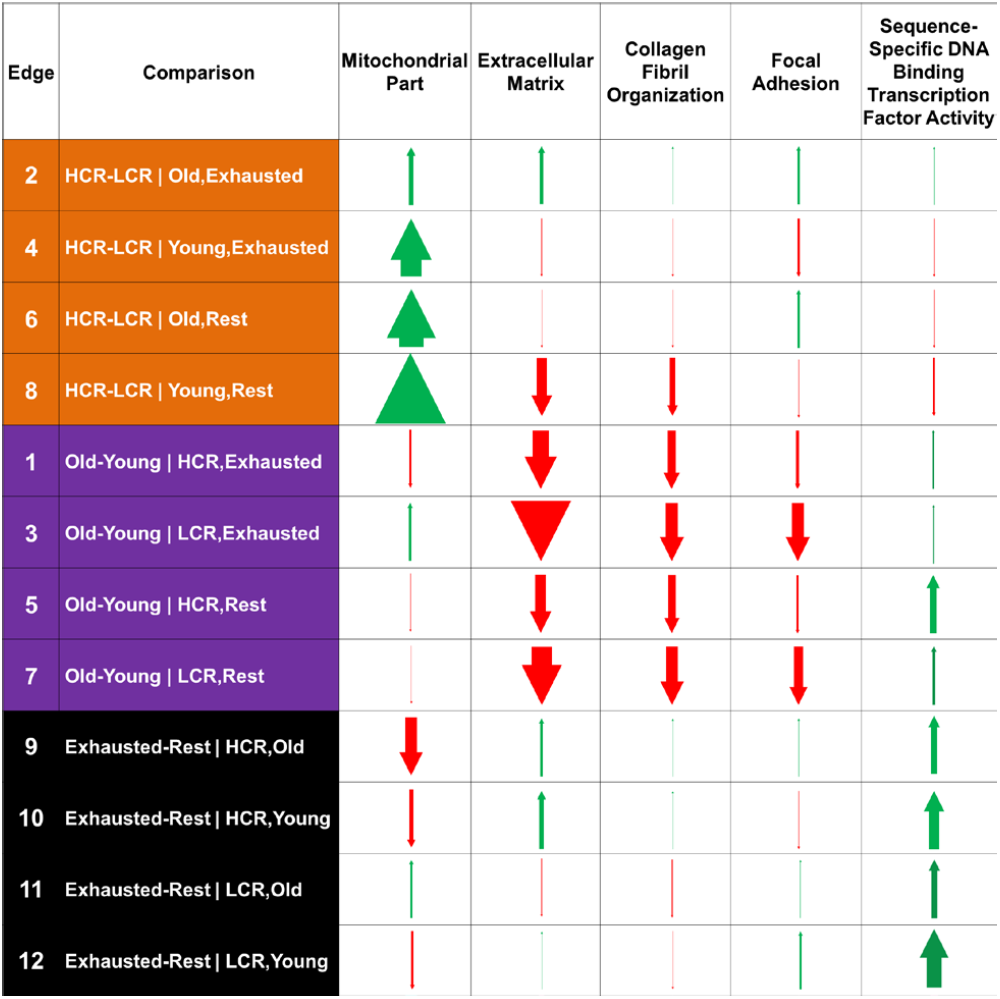
shown are the directions of the major pathway groups in our analysis. Green arrows denote that the pathway was up-regulated for the control group compared to the test group in each analysis, and vice versa for red arrows (control groups are HCR, old, and exhaust; test groups are LCR, young, and rest). The thickness of the arrow indicates the significance of the pathway for the respective analysis; with a thicker arrow representing smaller p-value.

**Figure 4.**
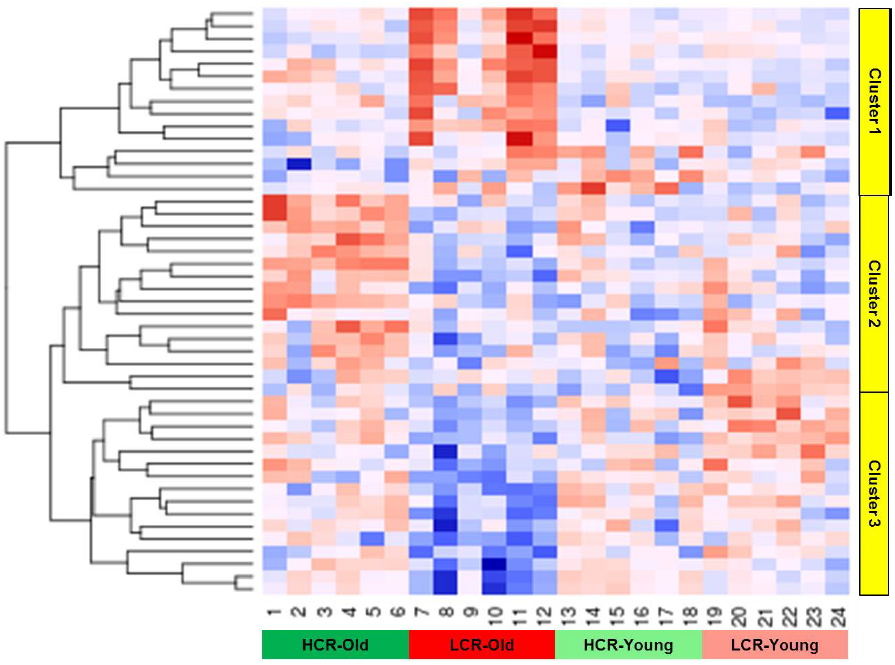
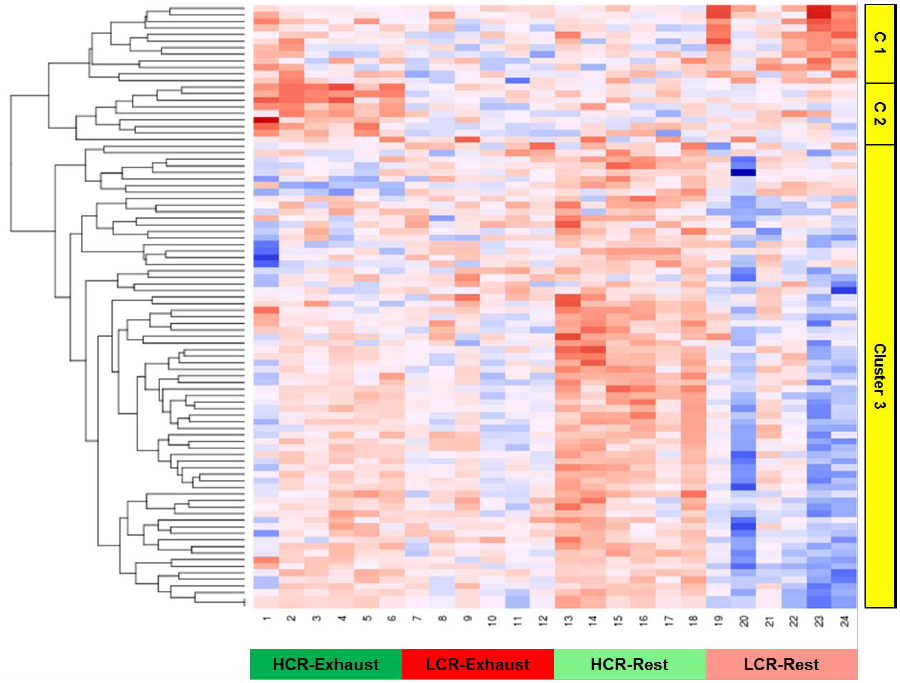
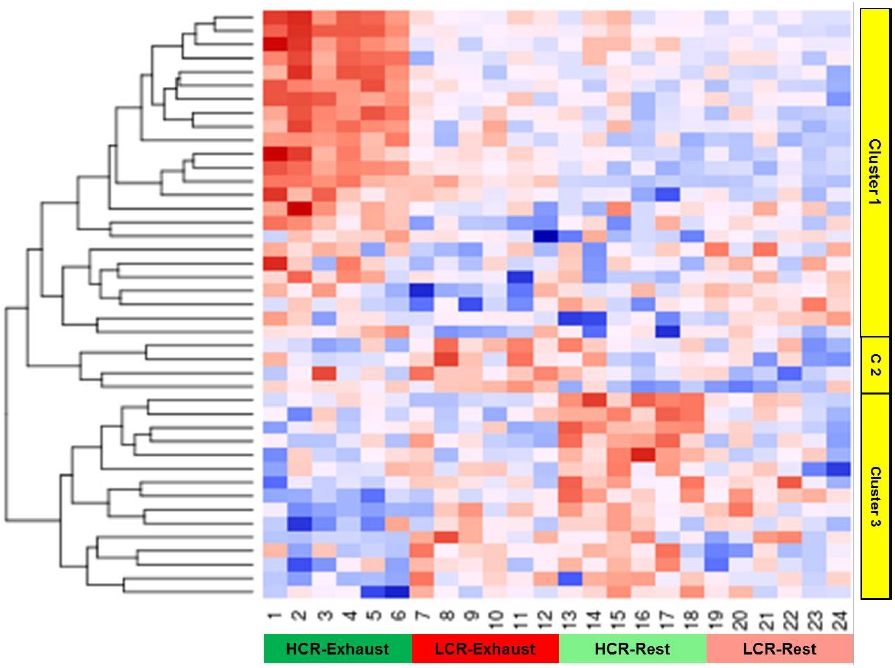
(A) shown is the heatmap of the 47 significant genes in the Muscle Structure Development pathway for the line-age interaction analysis for animals at Rest (bottom face). Each row shows the expression of a single gene across all samples, and the 24 samples at Rest are ordered by group (shown at the bottom) across the columns. The genes are grouped into clusters based on hierarchical cluster (shown on the right). The heatmap is colored in a blue-red spectrum, with low expression shown in dark blue and high expression shown in dark red. (B) shown is the heatmap of the 92 significant genes in the Mitochondrial Part pathway for the line-exercise interaction analysis for Old animals (left face). (C) shown is the heatmap of the 43 significant genes in Response to Biotic Stimulus pathway for the line-exercise interaction analysis for Young animals (right face).

For the line effect, HCR consistently shows up-regulated *Mitochondria Part* pathway compared to LCR (Edges-2, 4, 6 and 8) (**Figure 3a**).

For the age effect, old rats consistently show down-regulation in *Extracellular Matrix, Collagen Fibril Organization*, and *Focal Adhesion* pathways compared to young (Edges-1, 3, 5, 7) (**Figure 3b**). Further, aging in LCR (sides 3 and 7) results in more significant enrichment for all three pathways compared to HCR (sides 1 and 5) (**Table 2**).

For the exercise effect, exhausted rats consistently show up-regulated *Sequence-Specific DNA Binding Transcription Factor Activity* pathway compared to rats at rest (Edges-9, 10, 11, 12) (**Figure 3c**).

### Interaction effects

For each face of the cube, we assessed the interaction between two factors while keeping one factor constant (front=LCR; back=HCR; top=Exhausted; bottom=Rest; left=Old; right=Young), for a total of six analyses. We focused on the three faces with the most significant interactions (bottom: line-age for Rest; left: line-exercise for Old; right: line-exercise for Young) given the results from the main effect analysis.

For the bottom face, we assessed the interaction between line and age for only rats at Rest. The most significantly enriched pathway from our unidirectional *LRpath* analysis was *Muscle Structure Development* (p-value=1.1×10^-8^), which includes 248 total genes, out of which we found 47 to be below the nominal significance threshold (p<0.05). We analyzed these genes using Database for Annotation, Visualization and Integrated Discovery (*DAVID*) and found that these significant genes were most enriched for the *Muscle Cell Differentiation pathway* (8.4×10^-25^) [10]. The expression of these 47 genes formed three distinct clusters based on unsupervised hierarchical clustering for the 24 Rest samples (**Figure 4a**). We determined the pathway enrichment of each gene cluster using *DAVID*, and found that cluster 1 (15 genes) is most enriched for *Muscle Tissue Development* (p-value=3.5×10^-11^), cluster 2 (16 genes) is most enriched for Muscle Cell Differentiation (p-value=1.2×10^-9^), and cluster 3 (16 genes) is most enriched for *Muscle Organ Development* (p-value=1.9×10^-8^). Thus, we can see that the interaction effect between line and age for rats at Rest are largely due to muscle development pathways.

For the left face, we assessed the interaction between line and exercise for only Old rats. The most significantly enriched pathway from our unidirectional *LRpath* analysis was *Mitochondrial Part* (p-value=2.3×10^-6^), which includes 524 total genes, out of which we found 92 to be below the nominal significance threshold (p<0.05). When we analyzed these using *DAVID*, we found that these significant genes were still most enriched for the *Mitchondrial Part* pathway (4.6×10^-74^). As before, we determined the pathway enrichment of each gene cluster, and found that clusters 1 (12 genes) and 2 (9 genes) are both most enriched for *Mitchondrial Part* (p-value=1.1×10^-9^ and 1.4×10^-6^, respectively), and cluster 3 (71 genes) is most enriched for *Mitochondrion* (p-value=1.4×10^-57^), with *Mitochondrial Part* as a close second (p-value=6.1×10^-57^) (**Figure 4b**). Thus, we can see that the interaction effect between line and exercise for Old rats are largely due to mitochondrial pathways.

For the right face, we assessed the interaction between line and exercise for only Young rats. Given that the top two most significantly enriched pathways did not include large enough number of genes, we focused on the third most significantly enriched pathway, *Response to Biotic Stimulus* (p-value=4.4×10^-6^), which includes 342 total genes, out of which we found 43 to be below the nominal significance threshold (p<0.05). Using *DAVID*, we found that these significant genes were most enriched for the *Response to Bacterium* pathway (2.1×10^-11^). Cluster 1 (24 genes) is most enriched for *Response to Bacterium* pathway (p-value=1.5×10^-5^), cluster 2 (4 genes) is most enriched for *Response to Organic Substance* (p-value=5.9×10^-3^), and cluster 3 (15 genes) is most enriched for *Lymphocyte Proliferation* (p-value=1.4×10^-5^) (**Figure 4c**). We can see no clear primary driving pathway for the interaction effect between line and exercise for Young rats.

## Discussion

The ability to study three main effects (genetic background, exercise, aging) jointly and uncover interactions between them in a rat model is novel to our study. For all rats, the HCR-LCR difference is driven by mitochondrial pathways during both exercise and rest (**Table 1**). GO Biological Processes and KEGG Pathways show that genes associated with mitochondria are associated with substrate metabolism and oxidative metabolism, including Branched Chain Amino Acid metabolism, Fatty Acid metabolism, Oxidative Phosphorylation and Tricarboxylic Acid cycle (TCA) metabolism, are significantly higher in HCR muscle, as has previously been described [11]. We have recently found that the up-regulation of these pathways increases the capacity for non-glucose fuel utilization, delaying the ‘lactate threshold’ associated with glycolysis-mediated ATP production (Overmyer et al., in revision).

When the rats are at rest, the HCR-LCR gene expression difference is far greater for the old rats compared to the young (**Fig. 3a**), which could reflect a slowing of aging in HCR and the resulting longevity differences between HCR and LCR [12]. Interesting, this age-dependent difference disappears for the exhausted rats. This difference may be driven by genes in the mitochondria pathway; the difference between HCR-LCR is the strongest for young rats at rest, perhaps being driven by their intrinsic expression differences for mitochondrial genes (**Table 1**). The weaker HCR-LCR difference at exhaustion may be due to the up-regulation of the same exercise-related gene sets for both genetic backgrounds. The exhausted-rest difference is driven by the transcription activity pathway; with exhausted animals showing up-regulated transcription activity.

When we analyze the exhaustion-rest rats, we see that the difference is greatest for the young HCRs, and weakest for young LCRs (**Fig. 3b**). The associated genes are largely transcription factors and cofactors that are associated with muscle development or known to be responses to exercise which we have previously observed in trained human muscles [15]. These gene expression differences in HCR and LCR may simply be attributable to exercise duration, as HCRs run 9-fold longer distance that LCRs [13]. The old-young difference is driven by the ECM, collagen, and adhesion (aging indicator) pathways in both HCRs and LCRs during both exercise and rest (**Table 1**). When the rats are at rest, the old-young difference is far more prominent for LCRs compared to HCRs, showing that LCRs show greater magnitude of differential expression due to aging (**Fig. 3c**). In addition, age-effects in the aging indicator pathways are more pronounced in the LCRs compared to the HCRs. Both of these findings suggest that LCR’s lower innate aerobic capacity underlies faster aging.

As mammals age, the amount of collagen in our muscle increases, which results in muscle stiffness and reduced whole muscle function [14]. The interrelated pathways are responsible for the structural organization of muscle/matrix interactions. Collagen isotypes and other extracellular matrix components interact with Focal Adhesion Complexes containing integrin transduces signals associated with contractile forces in adult skeletal muscle [15, 16]. Focal Adhesion Kinase phosphorylation is activated by muscle contraction-induced association with integrins and transduces hypertrophy-related signaling [16] and can increase proteins associated with mitochondrial oxidation [17]. In addition to a greater decline in these pathways in the LCR with aging, The *Extracellular Matrix, Collagen Fibril Organization* and *Focal Adhesion* pathways also are significantly higher in HCR compared to LCR in Line-Effect, suggesting an interaction between the signaling pathways and alterations in muscle oxidation. Our analysis is interesting in that the collagen-related pathways are down-regulated in the old rats compared to the young rats. This finding is similar to those from a previous meta-analysis across humans and rodents for gene expression data among aging groups, in which the researchers also found collagen gene sets to be under-expressed with age, and explain it by reduced collagen deposition with aging [18].

The ability to study the effects of exercise to exhaustion is novel to our study. Physical exercise is a stressful event for all higher organisms; during which the host must recruit a series of physiological and morphological adaptations in order to achieve, one of the most important being muscle contraction. To sustain muscle contraction during exercise, the demand for adenosine triphosphate (ATP) can increase 1,000-fold compared to the resting state [19]. In our study, we found that exercise-induced expression difference (exhausted vs rest) for both old and young rats is greater in HCR (sides 9 and 10) than in LCR (sides 11 and 12). Our analysis of metabolite changes in HCR and LCR suggests a similar pathway to exhaustions, that is a depletion of glycogen which corresponds to the ‘lactate threshold’. This delay in exhaustion in the HCR would provide a more sustained signal to alter gene expression.

The broad premise that oxidative energy metabolism is mechanistically connected with longevity is attractive because it has the power to shape the multiplicity of biological networks that influence essentially every phenotype across a lifespan [20]. Additionally, endurance capacity fulfills the fundamental criteria for service as a biomarker of aging as suggested by The American Federation for Aging Research. That is, endurance capacity predicts the rate of aging accurately, represents a basic underlying process, can be tested repeatedly without harm, and can be evaluated in animals [21].

## Materials and Methods

### Ethics statement

This study was approved by the University Committee on Use and Care of Animals, Ann Arbor, Michigan (Approval Numbers: #08905 and #03797). The proposed animal use procedures are in compliance with University guidelines, and State and Federal regulations.

### Animals

We used old (~84-93 weeks of age) and young (~12-20 weeks of age) rats from HCR and LCR generations 29 and 32, respectively (**Supplementary Figure 2**). The study included eight groups; HCR-Old-Exhausted (H-O-E, n=6), HCR-Old-Rest (H-O-R, n=6), HCR-Young-Exhausted (H-Y-E, n=6), HCR-Young -Rest (H-Y-R, n=6), LCR-Old-Exhausted (L-O-E, n=6), LCR-Old-Rest (L-O-R, n=6), LCR-Young-Exhausted (L-Y-E, n=6), and LCR-Young -Rest (L-Y-R, n=6). For the exhausted rats, dissections were performed within 10 min after the maximal running distance was reached.

### Tissue and RNA extraction

We extracted skeletal muscle RNA from a total of 48 female animals (n=6 in each of the 8 group). Skeletal muscle tissue was obtained from the Extensor digitorum longus (EDL). All rats were dissected immediately after sacrificing, and all tissue samples were immediately weighed and snap frozen in liquid nitrogen, and stored at -80C. Total RNA was extracted from frozen tissue with a Trizol reagent (Invitrogen) and purified with an RNAse kit (Ambion).

### Gene expression microarray

We ran the skeletal muscle RNA on the Affymetrix Rat Gene ST 2.1 array. The microarray hybridizations were performed by the DNA sequencing core at the University of Michigan according to the manufacturer’s instructions. We used the Affymetrix Expression Console^™^ software to generate gene expression values from individual probe intensity (CEL) files. The microarray yielded a total of 19,607 transcript expressions.

### Gene expression data analysis

Data analysis were performed with an R software environment for statistical computing [22]. The data were normalized with quantiles normalization. Principal component analysis was performed using the *prcomp* function. The Euclidean distance between clusters was calculated with the *dist* function using the median expression values of each gene. Statistically significant differences in gene expression between test groups were tested using multiple regression with the *lm* function.

For the main effect analyses, we implemented the regression models:

exp ~ line + age + exercise
exp ~ line (for old and exhausted; edge 2)
exp ~ line (for young and exhausted; edge 4)
exp ~ line (for old and rest; edge 6)
exp ~ line (for young and rest; edge 8)
exp ~ age (for HCR and exhausted; edge 1)
exp ~ age (for LCR and exhausted; edge 3)
exp ~ age (for HCR and rest; edge 5)
exp ~ age (for LCR and rest; edge 7)
exp ~ exercise (for HCR and old; edge 9)
exp ~ exercise (for HCR and young; edge10)
exp ~ exercise (for LCR and old; edge 11)
exp ~ exercise (for LCR and young; edge 12)

For the interaction effect analysis, we implemented the regression models:

exp ~ line + age + line*age (for exhausted; top)
exp ~ line + age + line*age (for rest; bottom)
exp ~ line + exercise + line*exercise (for old; left)
exp ~ line + exercise + line*exercise (for young; right)
exp ~ age + exercise + age*exercise (for HCR; back)
exp ~ age + exercise + age*exercise (for LCR; front)

### Pathway analysis

Pathway enrichment was performed using *LRpath* [23] for all ~20K genes, p-values, and fold-change for each of the 12 comparisons. We tested for enrichment of Gene Ontology (GO) terms and Kyoto Encyclopaedia of Gens and Genomes (KEGG) pathways for the rat. *LRpath* allows us to perform both unidirectional and directional analyses; for unidirectional analysis, *LRpath* tests for gene sets that have significantly higher significance values than expected at random given a set of genes and p-values; for directional analysis, *LRpath* tests up-and down-regulated genes simultaneously given a set of genes, p-values, and fold-change between test groups to distinguish between up and down regulated gene groups.

---

**Supplementary Figure 1.**
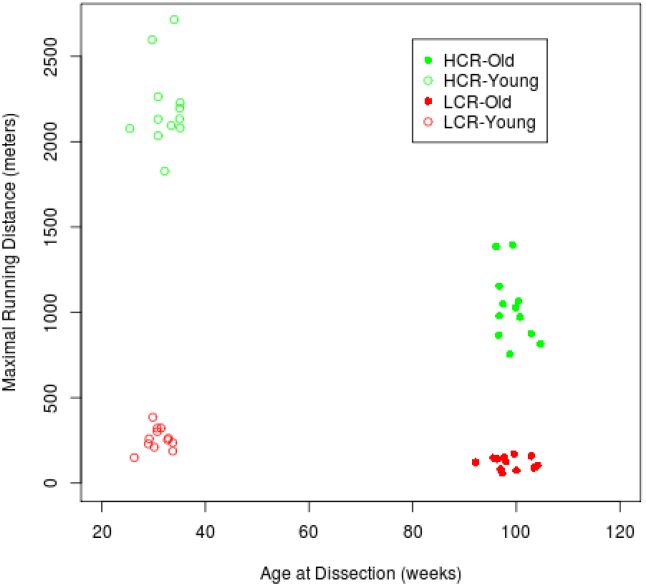
The scatterplot of age at dissection versus runner distance for the four groups. The samples are separated into four clusters corresponding to each group.

**Supplementary Figure 2.**
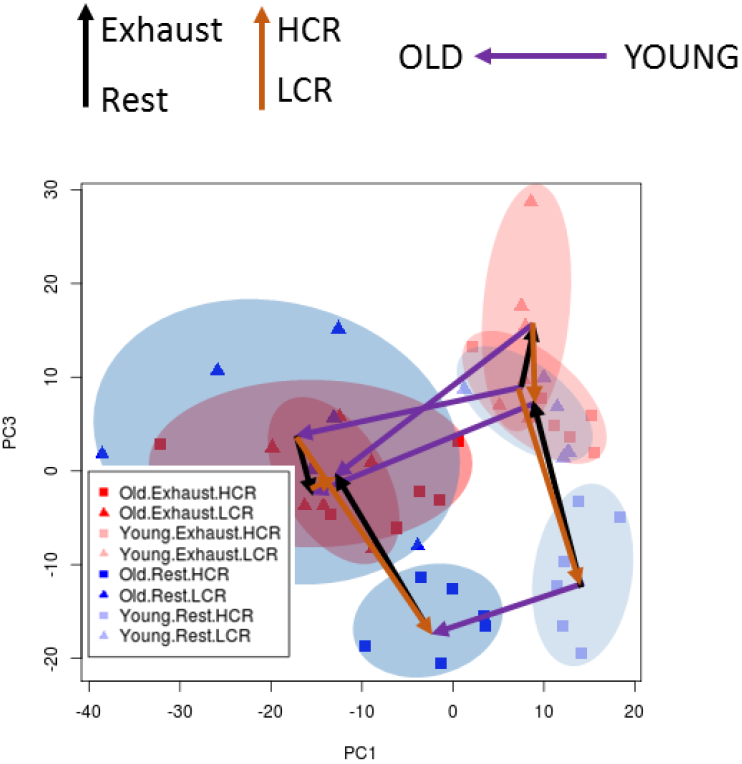
shown is the PC1 vs PC3 plot for 48 rats across the expression of ~20K transcripts. The samples are divided across 8 different groups in a 2×2×2 design to compare HCR and LCR, aged and young, and between rest and exhaustion (n=6 each). Groups are highlighted using different colored ovals to show clustering, and HCR-LCR are shown as squares and triangles, respectively. All of the exhausted animals are red, and animals at rest are blue. Old animals are shown as dark red/blue, and young animals are shown as light red/blue. Arrows are drawn from each cluster to show the direction of each group variable; purple arrows show direction of old-young animals; orange arrows show direction of HCR-LCR; black arrows show direction of exhaust-rest.

**Supplementary Table 1** – Tab-delimited spreadsheet with 19,607 rows (genes) and 31 columns for the collective regression results of the 15 main effect analyses (line, age, exercise, and edges 1-12). Column 1 is gene symbol, followed by 2 columns for each analysis (p-value, fold-change).

**Supplementary Table 2** – Tab-delimited spreadsheet with 4,146 rows (concepts) and 67 columns for the collective LRpath results of the 15 main effect analyses (line, age, exercise, and edges 1-12). Column 1 is concept name, column 2 is concept type, column 3 is number of genes in concept, followed by 4 columns for each analysis (odds ratio, p-value, FDR, and direction).

